# CD271 restrains the B1b cell antibody response in a T cell dependent manner

**DOI:** 10.1101/2024.01.08.574733

**Authors:** Marie-Ève Lebel, Carole A. Bonkoungou, Jade Thibault, Joyce Rauch, Kishore R. Alugupalli, Heather J. Melichar

**Affiliations:** Maisonneuve-Rosemont Hospital Research Center, Montreal, QC, Canada; Département de Microbiologie, Infectiologie et Immunologie, Université de Montréal, Montreal, QC, Canada; Research Institute of the McGill University Health Centre, Division of Rheumatology, Department of Medicine, McGill University, Montreal, QC, Canada; Department of Microbiology and Immunology, Sidney Kimmel Medical College, Thomas Jefferson University, Philadelphia, PA, United States; Sidney Kimmel Cancer Center, Thomas Jefferson University, Philadelphia, PA, United States; Département de médecine, Université de Montréal, Montreal, QC, Canada; Rosalind and Morris Goodman Cancer Institute, Department of Microbiology & Immunology, McGill University, Montreal, QC, Canada

## Abstract

Co-signaling molecules modulate T cell function and are essential to control the duration and amplitude of immune responses. These molecules belong to the immunoglobulin and tumor necrosis factor receptor (TNFR) superfamilies and have been extensively studied; however, the immunomodulatory role of several family members remains unknown. We show that CD271 (also known as p75 or NGFR), a TNFR superfamily member, is highly expressed on peritoneal B1 B cells, but not on conventional (B2) B cells at steady state. CD271 expression by B cells specifically restrains the response to T cell-independent type 2 antigens (TI-2). B1 cell-expressed CD271 maintains peritoneal CD4^+^ T cell quiescence, and augmented antibody responses to TI-2 antigens in CD271-deficient mice are dependent on CD4^+^ T cells. This increase in antibody production correlates with improved bacterial neutralization *in vitro* and survival after bacterial challenge *in vivo*. Further, our results suggest that CD271 can directly inhibit mouse and human T cell activity to the same extent as PD-L1, a known, negative regulator of T cell function. These results establish CD271 as a co-inhibitory molecule that could be targeted to improve the efficacy of vaccines against pathogenic encapsulated bacteria and to broadly modulate the T cell response in therapeutic contexts.

**eTOC summary:** Lebel et al. identify an unexpected role for CD271, a neurotrophin receptor, in directly inhibiting T cell function. B1 cell-expressed CD271 maintains CD4^+^ T cell quiescence; in the absence of CD271, T cells bolster antibody production to T-independent antigens, enhancing the response to encapsulated bacteria.

## Introduction

Co-signaling molecules are important regulators of lymphocyte activity that are essential to control the duration and amplitude of the immune response; they finely tune T cell activity to efficiently eliminate pathogens while preventing the development of immunopathology and autoimmune diseases^1^. Moreover, they have been the target of several therapeutic applications aimed at modulating the immune response in different contexts^2–4^. These cell surface proteins belong to the immunoglobulin and tumor necrosis factor receptor superfamilies (IgSF and TNFRSF, respectively) and have been intensely studied, but the immunoregulatory functions and relevant binding partners for several IgSF and TNFRSF members remain unknown. Of these, CD271 (also known as the low-affinity nerve growth factor receptor [NGFR], the p75 neurotrophin receptor [NTR], and TNFR super family member 16 [TNFSF16]) is particularly intriguing as it is reportedly expressed on multiple immunomodulatory cell types that include subsets of B cells, dendritic cells (DC), fibroblastic reticular cells, and mesenchymal stromal cells^5–12^. Importantly, aberrant CD271 expression and signaling is associated with dysregulated immune responses and human disease^8,10,13–15^.

CD271 is an orphan TNFR that does not bind to conventional ligands, and its role in modulating the T cell response has not been extensively studied. It is the low affinity receptor for all neurotrophins and plays an important role in cell survival and differentiation^16^. Thus, its role in immune cell modulation is generally considered in a cell-intrinsic manner with CD271 expression on immunomodulatory cells indirectly affecting T cell function. For example, lipopolysaccharide (LPS) exposure induces both nerve growth factor and CD271 expression on DCs, which boosts co-stimulatory molecule expression and inflammatory cytokine production to modulate T cell priming^5,13,17^; this, in turn, can impact T cell activity. In addition, there is evidence that inflammation induces CD271 expression, and that melanoma cells may escape T cell killing via dedifferentiation and subsequent down-regulation of tumor antigens upon autocrine signaling between CD271 and melanoma-produced neurotrophin^18,19^. Notably, however, several TNFRs not only signal in a cell-intrinsic manner, but they can also serve as ‘ligands’ that induce bidirectional or reverse signals through an interacting partner on other cells, in trans, to modulate the immune response^20–24^. It has been reported that CD271 binds to cell surface proteins such as human CD80, an important co-signaling molecule that belongs to the Ig superfamily^25–29^. The physiological relevance of these interactions is not known, and direct modulation of the T cell response by CD271 has not been considered.

RNA-seq data from the ImmGen database suggest that the strongest expression of *Ngfr* among immune cell populations is within the peritoneal B1b cell population^30^. B1b cells are key players in the establishment of protective immunity to encapsulated bacteria such as *Streptococcus pneumoniae* and *Salmonella* Typhi^31^. Unlike their conventional B2 cell counterparts that recognize protein antigens and require antigen-specific T cell help for antibody production, B1b cells produce antibodies following B cell receptor crosslinking by multivalent repetitive antigens like polysaccharides. As these antigens are not presented by MHC-II, they do not trigger antigen-specific T cell help and are called T-independent type 2 antigens (TI-2). However, non-cognate T cell help influences the antibody repertoire and IgG isotype usage of B1 cells^32–34^. Despite recent advances in our understanding of the regulation of the humoral response to TI-2 antigens, it is still challenging to develop efficient vaccines against multiple, life-threatening bacterial infections. A better understanding of the mechanisms that control the humoral response against TI-2 antigens is necessary to reduce morbidity and mortality caused by pathogenic encapsulated bacteria. Thus, we sought to determine the role of CD271 expression by B1b cells in the regulation of antibody production.

In this study, we show that CD271 is expressed by B1 cells isolated from the peritoneal cavity of wild type (WT) mice and demonstrate that B cell-specific CD271 expression restrains antibody production after immunization with TI-2 antigens in a CD4^+^ T cell-dependent manner. This increase in antibody production in the absence of B cell expressed CD271 has an important impact on the control of bacterial infection. In addition, we show that the extracellular domain of CD271 directly inhibits activation of mouse and human T cells *in vitro*. Given its expression on multiple immunomodulatory subsets, these results highlight the potential of CD271 to directly regulate the T cell response in various settings including autoimmunity and tumor control.

## Results

### CD271 is expressed by peritoneal B1 cells and restrains antibody production after NP-Ficoll immunization

To characterize the role of CD271 in the regulation of the humoral response, we first assessed CD271 expression within the B cell subpopulations of the peritoneal cavity and the spleen of naive WT mice. Interestingly, peritoneal B1a and B1b cells, but not B2 cells, express CD271 (Fig. 1A and Fig. S1A). In contrast, we did not detect CD271 on the surface of splenic B cell subsets at steady state (Fig. S1B and C). Thus, we sought to determine if CD271 expression impacts the humoral response to antigen using WT and germline CD271 knock-out (KO) mice. We took advantage of the well-known hapten 4-Hydroxy-3-nitrophenylacetic (NP) conjugated to different immunogens and the distinct effects of these immunogens on B cell responses. Specifically, NP coupled to LPS forms a TI-1 antigen that favors IgG2 and IgG3 antibody production by B1a cells while NP coupled to a polysaccharide, such as Ficoll, forms a TI-2 antigen that typically elicits NP-specific IgG3 antibody production predominantly by B1b cells^35^. In contrast, NP coupled to a protein carrier such as chicken gamma globulin (CGG) results in a T-dependent antigen primarily inducing IgG1 antibody production by B2 cells with the help of CD4^+^ T cells^35^. WT and CD271 KO mice produce similar levels of NP-specific IgG1 and IgG3 after intraperitoneal (i.p.) NP-CGG or NP-LPS immunization (Fig. 1B and C). Interestingly, however, mice lacking CD271 produce more NP-specific IgG3 after i.p. NP-Ficoll immunization compared to WT littermate controls (Fig. 1D). In addition, a higher number of NP-specific IgG3 antibody-secreting cells (ASCs) were observed in the mesenteric lymph nodes and the spleen of CD271 KO mice after NP-Ficoll injection, suggesting that CD271 limits the differentiation of ASCs and thus the amount of antibody produced after NP-Ficoll immunization (Fig. 1E-G). Importantly, no differences in total serum IgM and IgG antibodies, and IgG3, specifically, are detected in WT and CD271 KO mice at steady-state (Fig. S1D). We did, however, detect low to moderate levels of serum autoantibodies to double stranded DNA (dsDNA) and the phospholipid cardiolipin (CL) in a cohort of aged CD271 KO mice as compared to littermate controls, but autoantibodies to human β2 glycoprotein I (β2GPI) were negative in these mice (Fig. S1E).

**Figure 1.**
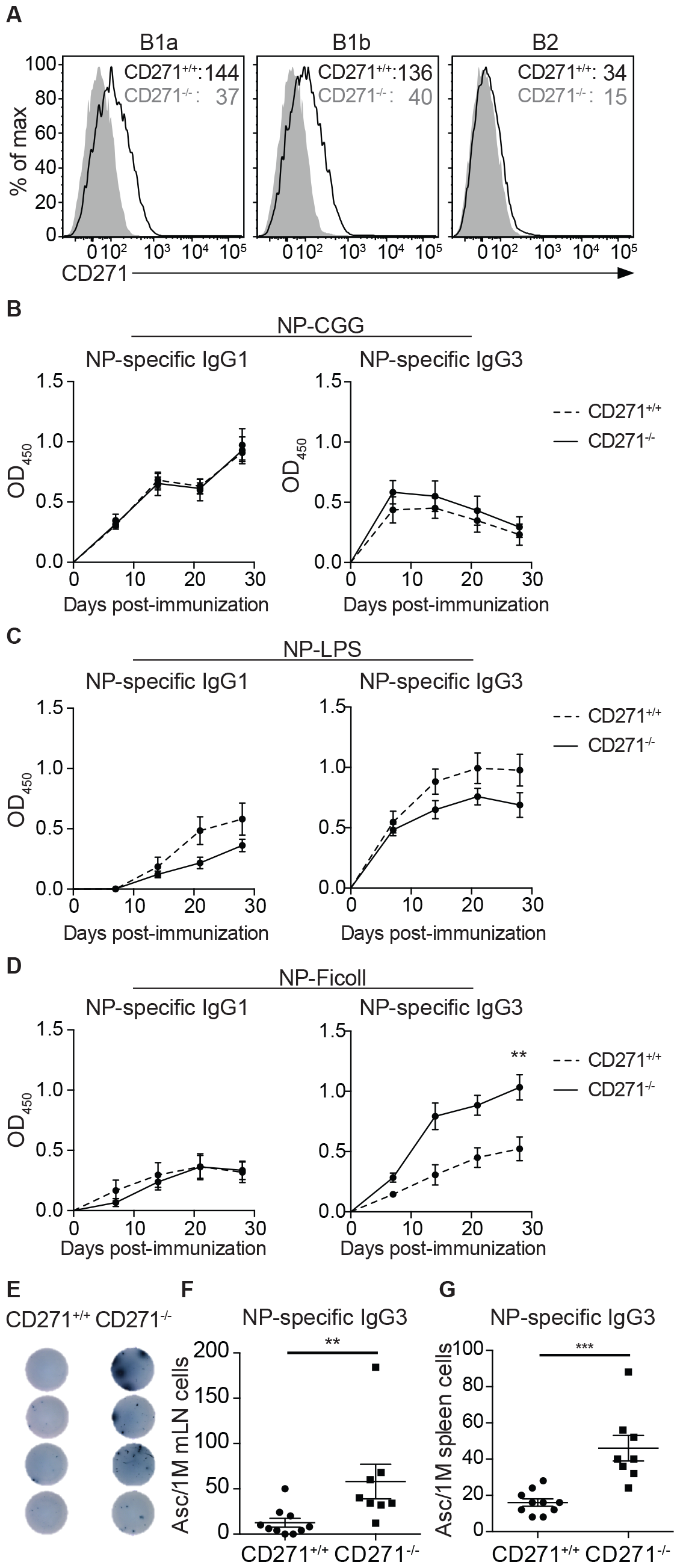
CD271 inhbits antibody production after NP-Ficoll immunization. (**A**) Representative histograms of CD271 expression on B cells isolated from the peritoneal cavity of CD271 WT (black line) or KO mice (gray histogram) of n=8 mice in each group. Numbers indicate median fluorescence intensity of CD271. WT (dashed line) or CD271 KO (solid line) mice were immunized with (**B**) NP-CGG, (**C**) NP-LPS, or (**D**) NP-Ficoll, and NP-specific antibodies in the serum were quantified by ELISA at 7, 14, 21, and 28 days post-immunization. Data are pooled from two or three independent experiments with 5-9 mice per group. Representative images (**E**) and compilation of the number of NP-specific IgG3 secreting cells measured at day 14 post-immunization in the mesenteric lymph nodes (**F**) and the spleen (**G**) of CD271 KO and WT littermate control mice by ELISPOT. Data are pooled from two independent experiments with 8-10 mice per group. Dots indicate individual mice. \**p* < 0.05, \*\**p* < 0.01, \*\*\**p* < 0.001

### CD271 expression by B cells restrains protection against TI-2 Ag-bearing pathogens

To confirm that CD271 expression by B cells themselves is important for the regulation of antibody production after NP-Ficoll immunization, CD19-cre^+/-^ (CD19-cre^+^) CD271^fl/fl^ mice (mice lacking CD271 specifically in B cells) were used^36,37^. We did not detect any significant difference in the level of total IgM, IgG3, or IgG2c antibodies in naive CD19-cre^+^ CD271^fl/fl^ mice as compared to CD19-cre^-^ CD271^fl/fl^ littermate controls (Fig. S2A). However, an increased production of NP-specific IgM, IgG3 and IgG2c was observed in these mice after i.p. NP-Ficoll immunization, suggesting that CD271 expression by B1 cells is sufficient to regulate Ig production in this context (Fig. 2A). As NP-Ficoll is an artificial TI-2 antigen, we sought to determine if CD271 also impacts antibody production after encounter with physiological pathogen-derived TI-2 antigens. To this end, B cell-specific CD271-deficient mice were i.p. immunized with *Salmonella enterica* serovar Typhi (S. Typhi) Vi polysaccharide (ViPS) or pneumococcal polysaccharide serotype 3 (PPS3)^38,39^. In both cases, Ag-specific IgG3 was higher in the absence of CD271 expression by B cells (Fig. 2B and C). Importantly, the higher ViPS-specific antibody titer correlates with a more robust bactericidal response against *S*. Typhi *in vitro* (Fig. 2D). Moreover, survival of PPS3-immunized B cell-specific CD271-deficient mice is improved after infection with a lethal dose of virulent *Streptococcus pneumoniae* (Fig. 2E). These results suggest that CD271 has a physiologically relevant immunoregulatory role in response to immunization with pathogen-derived TI-2 antigens.

**Figure 2.**
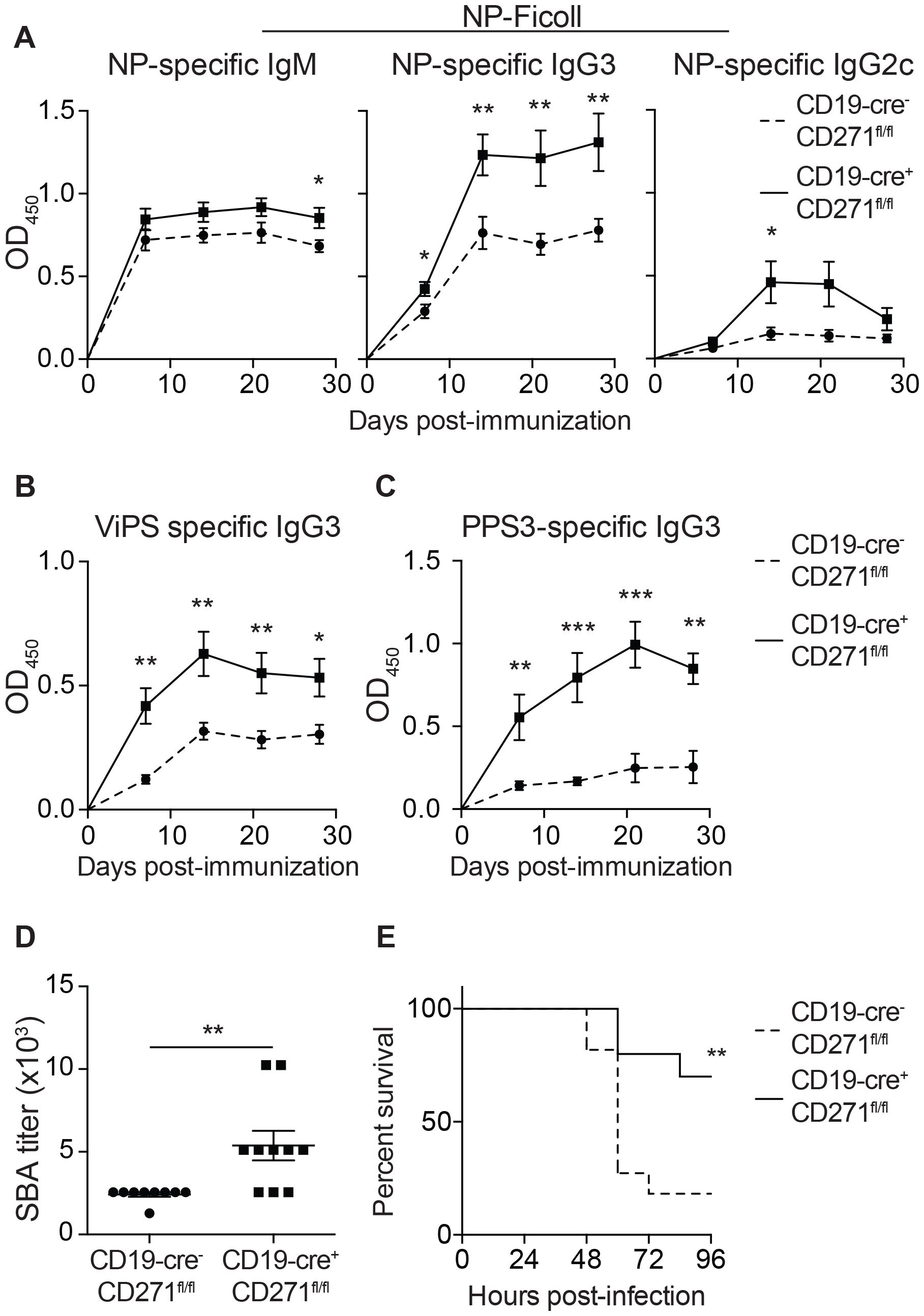
CD271 expression by B cells limits protection against TI-2 Ag-bearing pathogens. (**A**) CD19-Cre^+^CD271^fl/fl^ and littermate control (CD19-Cre^-^CD271^fl/fl^) mice were immunized with NP-Ficoll, and NP-specific IgM (left panel), IgG3 (middle panel), and IgG2c (right panel) in the serum were quantified by ELISA 7, 14, 21 and 28 days after immunization. Data are pooled from three independent experiments with 9-11 mice per group. CD19-Cre^+^CD271^fl/fl^ and littermate control mice were immunized with *Salmonella enterica* serovar Typhi Vi polysaccharide (ViPS) (**B**) or pneumococcal polysaccharide type 3 (PPS3) (**C**), and ViPS- or PPS3-specific IgG3 in the serum were quantified by ELISA 7, 14, 21, and 28 days post-immunization. Data are pooled from three independent experiments with 9-10 mice per group. (**D**) Serum bactericidal (SBA) titers against *S*. Typhi strain Ty2 were determined at 28 days post ViPS immunization. Data are pooled from two independent experiments with 8-10 mice per group. Dots indicate individual mice. (**E**) PPS3 immunized CD19Cre^+^CD271^fl/fl^ and littermate control mice were infected with 1.5 x 10^5^ CFU *Streptococcus pneumonia* WU2 i.p., and the survival of the mice was assessed. Data are pooled from two independent experiments with 10-11 mice per group \**p* < 0.05, \*\**p* < 0.01, \*\*\**p* < 0.001

### Dysregulated antibody production after TI-2 antigen immunization in the absence of CD271 depends on CD4^+^ T cell help

As B1b cells are the main B cell subpopulation responsible for IgG3 production after NP-Ficoll immunization^40^, we determined whether the percentage of B1b cells was higher in the peritoneal cavity of naive CD19-cre CD271^fl/fl^ mice as compared to other B cell populations. At steady state, the absence of CD271 expression on B cells does not affect the percentage of the different B cell subtypes in the peritoneal cavity (Fig. S2B). However, 28 days after NP-Ficoll immunization, there was a skewing of B cell subsets towards a higher percentage of B2 cells and a reduced percentage of B1a cells in the peritoneal cavity. Interestingly, the relative percentage of B1b cells remained similar between CD19-cre^+^ CD271^fl/fl^ mice and littermate controls (Fig. S2C).

Although CD4^+^ T cells are not required for antibody production following NP-Ficoll immunization, they can produce cytokines that augment isotype switching and recruit other types of B cells to the peritoneal cavity, thus influencing the humoral response against T cell-independent antigens^32,33^. We therefore evaluated the role of CD4^+^ T cells in this model. At steady state, there is a small, but significant increase in the activation of CD4^+^ T cell in the peritoneal cavity of CD19-cre^+^ CD271^fl/fl^ mice as assessed by IFN-γ production, as well as CD43 (1B11) and CD25 expression as compared to littermate controls (Fig. 3A-D). No difference in T cell activation was observed in the spleen of CD19-cre CD271^fl/fl^ mice compared to WT mice, consistent with the observation that CD271 is not significantly expressed by splenic B cells at steady state (Fig. S3). To specifically evaluate whether CD4^+^ T cells are involved in the elevated NP-specific IgG3 production observed after NP-Ficoll immunization in CD19-cre^+^ CD271^fl/fl^ mice, mice depleted of CD4^+^ T cells were immunized with NP-Ficoll. Strikingly, CD4^+^ T cell depletion prevented the increase in NP-specific antibody production, confirming the involvement of CD4^+^ T cells in this process (Fig. 3E).

**Figure 3.**
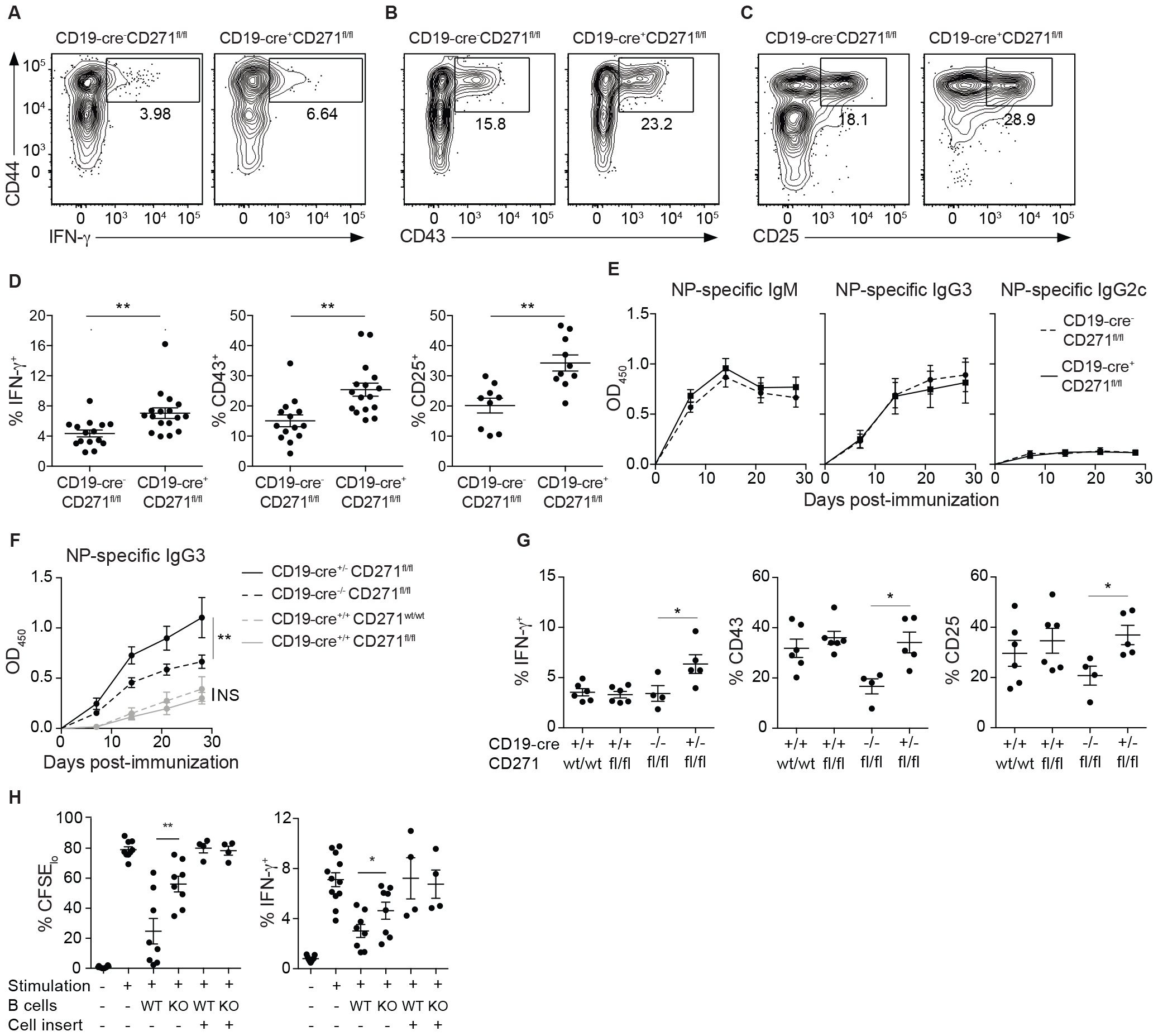
B1 cell-expressed CD271 inhibits the humoral response to NP-Ficoll in a CD4^+^ T cell-dependent manner. CD4^+^ T cells were isolated from the peritoneal cavity of naive CD19-Cre^+^CD271^fl/fl^ or littermate control (CD19-Cre^-^CD271^fl/fl^) mice and were stimulated with PMA/iono for 4 hours in the presence of Brefeldin A. Representative histograms (**A**-**C**) and compilation of the percentage (**D**) of IFN-Ψ^+^, CD43^+^, and CD25^+^ CD4^+^ T cells. Data are pooled from two to four independent experiments with 9-17 mice per group. Dots indicate individual mice. (**E**) CD4^+^ T cells were depleted from CD19-cre^+^ CD271^fl/fl^ and littermate control mice at day -1, 0 and 14 post-NP-Ficoll immunization, and NP-specific IgM, IgG3 and IgG2c were quantified by ELISA in the serum at day 7, 14, 21, and 28 post-immunization. Data are pooled from two independent experiments with 7 mice per group. (**F**) CD19 sufficient (CD19-cre^-/-^ or CD19-cre^+/-^) or deficient (CD19-cre^+/+^) mice lacking or not CD271 were immunized with NP-Ficoll, and NP specific IgG3 in the serum was quantified by ELISA 7, 14, 21 and 28 days after immunization. Data are pooled from two independent experiments with 5-9 mice per group. (**G**) CD4^+^ T cells were isolated from the peritoneal cavity of CD19 sufficient or deficient mice lacking or not CD271 and were stimulated with PMA/iono for 4 hours in the presence of Brefeldin A. Compilation of the percentage of IFN-γ producing, CD43^+^, and CD25^+^ CD4^+^ T cells. Data are pooled from two independent experiments with 4-6 mice per group. (**H**) Splenic T cells from WT mice were activated with anti-CD3/CD28 antibodies in the presence of B cells isolated from the peritoneal cavity of CD271 WT or KO mice for three days, together or separated by a cell culture insert, and were further stimulated with PMA/iono for four hours. Compilation of the proportion of CFSE_lo_ or IFN-γ producing CD4^+^ T cells. Data are pooled from two to four independent experiments with 4-12 mice per group. Dots indicate individual mice. \**p* < 0.05, \*\**p* < 0.01, \*\*\**p* < 0.001

B1a cells are known to have immunoregulatory properties that impact T cell function^41–43^. Therefore, we took advantage of the fact that CD19 KO mice lack B1a cells as well as marginal zone B cells (MZB) to asses the role of CD271 on these cell subsets^44,45^. In CD19-cre mice, the cre recombinase gene was knocked into the CD19 allele, and mice homozygous for the cre allele (CD19-cre^+/+^) lack endogenous CD19 expression^37^. We immunized CD19-cre^+/+^ mice and tested antibody production. In the absence of CD19 expression, the lack of CD271 expression by the remaining B cells does not lead to increased NP-specific IgG3 antibody production, suggesting that either B1a/MZB cells or CD19 itself is implicated in the elevated antibody production following NP-Ficoll immunization (Fig. 3F). The activation level of peritoneal CD4^+^ T cell was similarly unaffected in CD19 KO mice with or without CD271 expression in B cells (Fig. 3G), suggesting that CD271 expression by B1a cells may regulate peritoneal T cell quiescence.

To begin to tease apart the mechanism by which B1 cell-expressed CD271 modulates T cell activity, we performed co-culture assays with B cells harvested from the peritoneal cavitiy of WT or CD271 KO mice. Interestingly, when T cells are activated *in vitro* in the presence of B cells isolated from the peritoneal cavity, there is reduced CD4^+^ T cell proliferation and IFN-γ production in a cell-contact dependent manner. Further, CD271-deficient B cells are less immunosuppressive in this assay than their WT counterparts, suggesting that CD271 expression on peritoneal B1 cells might play a role in directly modulating T cell activity (Fig. 3H).

### The extracellular domain of CD271 directly inhibits mouse and human T cell activation *in vitro*

To determine whether CD271 can directly modulate T cell function, we took advantage of established *in vitro* activation assays. First, naive T cells were enriched from WT mouse spleen and activated with anti-CD3 and anti-CD28 antibodies with or without plate-bound recombinant proteins composed of the extracellular domain of CD271 or the well-known inhibitory ligand PD-L1 fused to the fragment crystallizable (Fc) region of IgG1 (CD271-Fc and PD-L1-Fc, respectively). When murine T cells are activated in the presence of CD271-Fc, T cell proliferation is inhibited as strongly as T cells activated in the presence of PD-L1-Fc (Fig. 4A and B). In addition, CD25 expression as well as IFN-γ production is as repressed by CD271-Fc as it is with PD-L1-Fc, suggesting a direct inhibition of T cell activity by CD271 (Fig. 4C-E). Moreover, CD271-Fc inhibited the activation and function of human peripheral blood T cells at levels similar to, or greater than, PD-L1-Fc (Fig. 4F-J). Together, these data suggest that CD271 directly inhibits T cell activity and highlight a possible broader impact of CD271 in the regulation of T cell responses.

**Figure 4.**
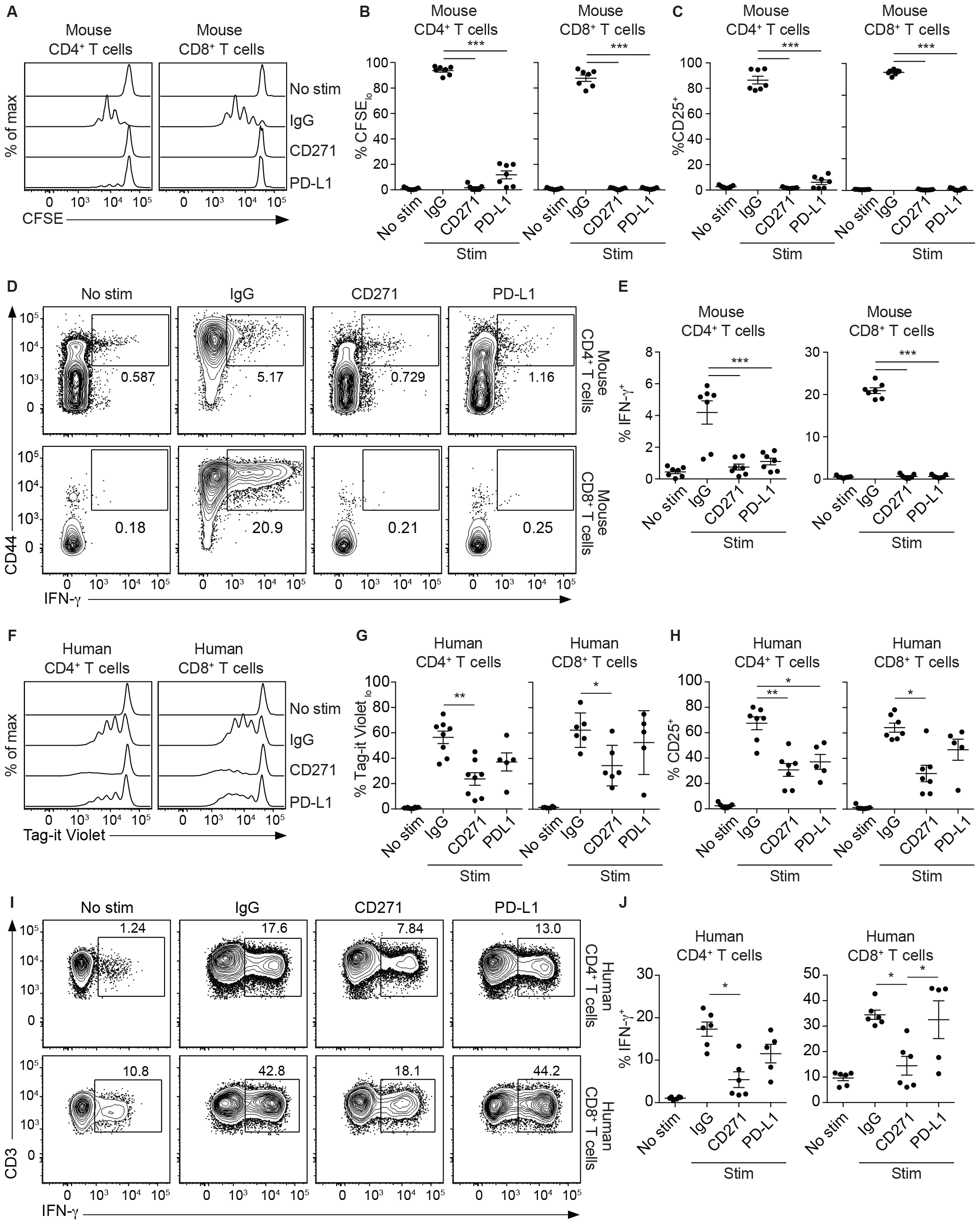
The extracellular domain of CD271 represses T cell activation. Naive murine T cells enriched from the spleen of WT mice were activated with anti-CD3 and anti-CD28 antibodies in the presence of different Fc-proteins. Cell proliferation and cytokine production were analyzed 3 days post-activation. Representative histograms of CFSE dilution (**A**) and compilation of the percentage of CFSE_lo_ CD4^+^ and CD8^+^ T cells (**B**). (**C**) Percentage of CD25^+^ CD4^+^ and CD8^+^ T cells 3 days post-activation. Representative flow plots (**D**) and compilation of the percentage (**E**) of IFN-γ producing CD4^+^ and CD8^+^ T cells 4 h after PMA/iono restimulation in the presence of Brefeldin A. Data are pooled from two independent experiments with 7 mice per group. (**G**) Human T cells enriched from PBMCs were activated with anti-CD3 and anti-CD28 antibodies in the presence of different Fc-proteins. Cell proliferation, cytokine production and cell activation were analyzed 4 days post-activation. Representative histograms of Tag-it Violet dilution (**F**) and compilation of the percentage of Tag-it Violet _low_ cells (**G**). (**H**) Percentage of CD25^+^ CD4^+^ and CD8^+^ T cells. Representative histograms (**I**) and compilation of the percentage of IFN-γ producing T cells (**J**) 4 h after PMA/iono restimulation in the presence of Brefeldin A. Data are pooled from two independent experiments with 5-6 donors per group. Dots indicate individual mice/human healthy donors. \**p* < 0.05, \*\**p* < 0.01, \*\*\**p* < 0.001

## Discussion

Encapsulated bacteria such as *Streptococcus pneumoniae, Neisseria meningitis, Haemophilus influenzae* and *Salmonella* Typhi remain public health challenges worldwide, especially in children. While protection against these pathogens is mediated by antibodies targeting their capsule polysaccharides, polysaccharide vaccines are poorly immunogenic in young children. Fortunately, the development of polysaccharide-protein conjugate vaccines was able to overcome this problem. Nevertheless, as the development of diseases caused by many encapsulated bacteria occurs very quickly, the presence of high titers of antibodies at the time of infection is crucial to provide adequate protection. In the present study, we demonstrate that CD271 expression by B cells restrains antibody production after immunization with TI-2 antigens (e.g., NP-Ficoll, PPS3, and ViP); B cell specific CD271-deficient mice show stronger antibody production. Importantly, this improved response leads to better bacterial neutralization *in vitro* as well as improved protection after *in vivo* challenge with a pathogenic encapsulated bacterium, suggesting that blocking CD271 during polysaccharide vaccine administration could lead to improved defense against diseases caused by pathogenic encapsulated bacteria.

Multiple mechanisms exist whereby CD271 might regulate the humoral response. CD271 is a co-signaling molecule and a neurotrophin receptor that could regulate cell survival, proliferation, and differentiation of the B1 cells intrinsically. Indeed, dysregulated neurotrophin signaling by B cells is associated with systemic lupus erythematosus (SLE)^10,11^. However, in the absence of CD271 expression by B cells, we did not detect any major difference in B cell numbers and phenotype at steady state. In addition, CD4^+^ T cells are more activated and produce more IFN-γ in absence of CD271 expression by B cells in naive mice, and we demonstrate that the improved humoral response to TI-2 antigen in the absence of CD271 expression by B cells is dependent on CD4^+^ T cells. We provide additional evidence that CD271 expression by B cells directly inhibits T cell activation and function *in vitro*. Thus, although we cannot exclude a cell intrinsic effect of CD271 expression by B1 cells, it is tempting to speculate that CD271 is an inhibitory ligand for a receptor on T cells.

B1 cells, and more specifically B1a cells, are known to have immunoregulatory properties that impact T cell function. For example, PD-L2 expression on the surface of B1a cells inhibits IL-5 production by T cells, thus impacting natural antibody production^41^. B1a cells can also convert naive CD4^+^ T cells into regulatory T cells producing high levels of IFN-γ and IL-10 in a CD86-dependent pathway. In addition, peritoneal B cells were shown to inhibit IFN-γ production by CD4^+^ T cells partly through an IL-10 dependent mechanism, which impacts colitis development^43^. Thus, it is possible that CD271 expression by B1a cells impacts the function of CD4^+^ T cells, which then influence the response of B1b cells to TI-2 antigen immunization.

Although the T-dependent humoral response could also benefit from an increased CD4^+^ T cell function, we did not observe any impact of the absence of CD271 on the production of antibodies following i.p. immunization with the T-dependent antigen NP-CGG. One possible explanation is that the T-dependent antigen response is preferentially mounted in the spleen, while CD271 expression on B1 cells was primarily observed in the peritoneal cavity. In line with this, splenic, but not peritoneal, CD4^+^ T cell activation and IFN-γ production at steady state was similar between control and B cell specific CD271-deficient mice, suggesting that the peritoneal microenvironment is important for the role of CD271 in the regulation of the humoral response. It is possible that some cytokines, such as IFN-γ (which is known to induce CD271 expression^18^), are present at a higher level in the peritoneal cavity as compared to the spleen, thus leading to this discrepancy. It will be interesting to explore whether CD271 also regulates antibody production to TI-2 antigens when the immunization route differs.

In conclusion, the key role of TNFRSF members in the fine tuning of the immune response is well recognized. However, the role of several superfamily members remains unclear. Here, we uncover a previously unappreciated role for CD271 in CD4^+^ T cell-dependent regulation of the humoral response to TI-2 antigens. The identification of CD271 as a potential direct regulator of T cell function and its expression by several immunoregulatory cell populations, including cancer cells, opens up a wealth of scenarios in which CD271 might act to modulate the T cell response.

## Methods

### Study Approval

Human peripheral blood was obtained from donors after written informed consent and in accordance with Research Ethics Committee guidelines at the Maisonneuve-Rosemont Hospital. All Research Ethics Committee guidelines respected the Declaration of Helsinki principles. All animal protocols were approved by the local Animal Care Committee at the Maisonneuve-Rosemont Hospital in line with the Canadian Council on Animal Care guidelines.

### PBMC isolation

Human peripheral blood was obtained from healthy donors. Peripheral blood mononuclear cells (PBMCs) were isolated by density gradient centrifugation over Ficoll-Paque (GE Healthcare) and cryopreserved until use. Cryopreserved PBMCs were thawed and incubated for 2 h at 37C before use.

### Mice

CD271 KO mice (B6.129S4-*Ngfr*^*tm1Jae*^/J, Stock# 002213) and CD19-cre (B6.129P2(C)-*Cd19*^*tm1(cre)Cgn*^/J Stock# 006785) mice were purchased from The Jackson Laboratory^37,46^. CD271 flox mice were kindly provided by Dr. Brian Pierchala (University of Michigan, current address Indiana University School of Medicine), and they were crossed with CD19-cre mice^36,37^; the breeding strategy ensured that the CD19-cre allele was not homozygous unless otherwise noted. Mice were maintained in a specific pathogen-free environment at the Maisonneuve-Rosemont Hospital Research Center. Both male and female mice were used unless indicated; no sex-dependent phenotypes were observed.

### Immunization and infection

Six-to twelve-week-old mice were immunized with 25 g 4-Hydroxy-3-nitrophenylacetic hapten conjugated to AminoEthylCarboxyMethyl-Ficoll (NP-Ficoll, LGC Biosearch Technologies), 50 µg NP-CGG (Chicken Gamma Globulin, LGC Biosearch Technologies) precipitated in alum (Cedarlane), 0.5 µg pneumococcal polysaccharide type 3 (PPS-3, ATCC, USA), or 2.5 µg Vi polysaccharide (ViPS, FDA) of *Salmonella enterica* serovar Typhi via i.p. injection. Mice were bled at day 7, 14, 21 and 28 post-immunization to quantify antigen-specific antibody production. In some cases, CD4^+^ T cells were depleted by injecting 250 µg anti-mouse CD4 (GK1.5, Leinco) i.p. at day -1, 0 and 14 of immunization. PPS-3 immunized mice were infected with 1.5 x 10^5^ CFU *S. pneumoniae* i.p. at > 28 days post-immunization. Mice were monitored every 12 h and euthanized upon demonstrating signs of morbidity.

### Serum bactericidal assay

A serum bactericidal assay was performed as previously described^47^. Log-phase culture of *S*. Typhi strain Ty2 was prepared in Luria–Bertani (LB) broth with 10 mM NaCl and was adjusted to 2.5-5.0 x 10^4^ CFU/ml in Dulbecco’s phosphate-buffered saline (PBS). Serum samples from ViPS-immunized, as well as unimmunized, mice were heat inactivated for 30 minutes at 56°C, and 2-fold serial dilutions in normal saline were performed in a round-bottom polypropylene 96-well plate. Ten microliters (250-500 CFUs) of *S*. typhi strain Ty2, 12.5 µl of baby rabbit complement (Pel-Freez), and 27.5 l of PBS were added to the 96-well plate containing 50 µl of serially diluted serum. The assay was performed in triplicate and incubated at 37°C for 120 min with gentle rocking. After incubation, 10 µl of the mix was plated on LB agar. Serum bactericidal antibody titers were defined as the reciprocal of the highest dilution that produced 50% killing in relation to negative control wells containing naive serum.

### ELISA

For quantification of antigen-specific antibodies, EIA/RIA 96-Well Plates (Corning) were coated overnight with 0.1 µg/ml NP-BSA in NaHCO_3_ 0.1 M pH 9.6, 10 µg/ml PPS3 in PBS, or 2 µg/ml ViPS in PBS. For total antibody quantification, plates were coated with 2 µg/ml anti-mouse-IgM or IgG (Biolegend) in NaHCO_3_ 0.1 M pH 9.6. Plates were blocked with PBS, containing 0.2% Tween-20 and, 1% BSA, for 1 h at room temperature and diluted sera were incubated for 1 h 30 at room temperature. Antigen-specific antibodies were detected using HRP-conjugated anti-mouse IgM, IgG3 or IgG2c (Jackson ImmunoResearch Inc.) and TMB-substrate (Biolegend). The reaction was stop by adding 2 N H_2_SO_4_, and the optical density was read at 450 nM. Antibodies to β2GPI, CL, and dsDNA were determined by ELISA on serum diluted 1/50 in PBS containing 0.3% gelatin and 10% fetal bovine serum, with serum from an MRL/MpJ-*Tnfrsf6 lpr* mouse, a β2GPI/LPS induced model of SLE, and a naive C57BL/6 mouse served as positive and negative internal controls for the ELISAs, as described previously^48,49^.

### ELISPOT

ELISPOT 96 well plates (MultiScreenHTS IP Filter Plate, 0.45 µm, Sigma-Aldrich) were precoated with 2 µg/ml NP-BSA in PBS over night at 4°C. The plates were washed and blocked with Roswell Park Memorial Institute medium (RPMI, Thermofisher Scientific) containing 10% fetal bovine serum (FBS, Thermofisher Scientific) for 2 h at 37°C. 1 x 10^6^ cells from the mesenteric lymph nodes in RPMI 10% FBS were cultured for 24 h at 37°C, 5% CO_2_. The plates were washed and incubated for 24 h at 4°C with HRP-conjugated anti-mouse IgG3 (Jackson ImmunoResearch Inc.) diluted 1/10000 in RPMI 10% FBS. Finally, the plates were washed, and the spots were revealed with TMB substrate (Mabtech, Inc.).

### *In vitro* T cell activation

Plates were coated with 1.25 µg/ml anti-CD3 (145-2C11 or OKT3, Biolegend) with or without 10 µg/ml Fc proteins (R&D Systems Inc.). Mouse T cells were enriched with EasySep™ Mouse T Cell Enrichment Kit (Miltenyi Biotec) according to manufacturer instructions and labeled with 5 M carboxyfluorescein succinimidyl ester (CFSE, Thermofisher Scientific) for 10 min at 37°C. Human T cells were enriched from PBMCs using the EasySep Negative Human T Cell kit (Miltenyi Biotec) according to manufacturer instructions and labeled with 5 M Tag-it Violet (Biolegend) for 20 minutes at 37°C. The staining was quenched with RPMI, containing 10% FBS for 10 min at 37°C. Mouse and human T cells were washed three times with RPMI 10% FBS and 2.5 x 10^5^ cells were added to each well with 2 µg/ml soluble anti-CD28 (37.51 or CD28.2, Biolegend). For co-culture experiments, cells were harvested from the peritoneal cavity of WT or CD271 KO mice, and B cells were enriched using anti-CD19-biotin (4B12, Biolegend) and the Mouse Streptavidin RapidSpheres Isolation kit (Stemcell Technologies). 1.25 x 10^6^ CFSE-labeled splenic T cells (enriched as above) were pre-incubated, or not, with 312,500 B cells for 1 h prior to adding to 24 well plates coated with 5 µg/ml anti-CD3 with or without a permeable polycarbonate cell culture insert (Fisher Scientific) in the presence of 2 µg/mL soluble anti-CD28. The cells were incubated for 72-96 h at 37°C, 5% CO_2_ before analysis.

### Flow cytometry

Flow cytometry analysis of mouse surface proteins was performed with antibodies against the following molecules: CD19 (6D5), CD4 (GK1.5), CD8 (53-6.7), CD23 (B3B4), CD25 (PC61), CD43 (1B11), CD44 (IM7), F4/80 (BM8), TCR (H57-597) (Biolegend), and CD5 (53-7.3) (BD Biosciences). CD271 expression was detected using anti-p75 NGF Receptor (polyclonal, Abcam) followed by anti-rabbit IgG (polyclonal, Biolegend). Flow cytometry analysis of human surface proteins was performed with the following antibodie: anti-CD3 (OKT3), -CD4 (OKT4), -CD8 (RPA-T8) and -CD25 (BC96). Staining was performed for 20 min at 4°C. For intracellular cytokine staining, cells were cultured with 50 ng/ml phorbol myristate acetate (PMA) and 1 µg/ml ionomycin (iono) in the presence of brefeldin A (10 µg/ml) for 4 h at 37°C. Following staining for surface antigens as described above, cells were stained with anti-IFN-γ antibody (XMG1.2 or 4S.B3) (Biolegend) using fixation/permeabilization buffers (BD Biosciences) according to the manufacturer’s instructions. For B cell analysis, Fc receptors were blocked using anti-CD16/32 (2.4G2) (Leinco) antibody, and the cells were stained with Zombie Aqua fixable viability dye (BioLegend) for 15 min at room temperature prior to cell surface staining. Flow cytometry analyses were performed on a BDLSR Fortessa X20 flow cytometer (BD Biosciences), and data were analyzed using the FlowJo software (BD Biosciences).

### Statistical analysis

Each experiment was repeated independently with similar results at least two times. For comparison between two groups, data were analyzed with the two-tailed Mann–Whitney U test. Data from more than two groups were analyzed with Kruskal–Wallis test followed by Dunn’s post-hoc comparisons, two-sided. All values where *p* < 0.05 were considered statistically significant. Analyses were performed with GraphPad Prism version 6. \**p* < 0.05, \*\**p* < 0.01, \*\*\**p* < 0.001.

## Data availability

The data are available from the corresponding author upon request. All the data needed to evaluate the conclusions of the paper are present in the paper or the online supplemental material.

## Acknowledgements

We are thankful for scientific discussion and constructive feedback from Ben Haley and Sébastien This as well as helpful review of the manuscript by Marilaine Fournier. We thank Martine Dupuis and Frédéric Duval for assistance with flow cytometry as well as the staff of the animal facilities for the maintenance of the mouse colonies at Maisonneuve-Rosemont Hospital Research Center. We appreciate technical support from Rebecca Subang. David Briles and W. Edward Swords kindly provided the *S. pneumoniae*. This work was supported by grants from the Canadian Cancer Society (703856), Cancer Research Society (24007), and the Canadian Institutes of Health Research (CIHR) (PJT-185949) awarded to H.J.M as well as CIHR grant (PJT-159652) to J.R. J.T. received a PREMIER summer fellowship from the Université de Montréal. H.J.M. is a senior scholar of the FRQS.

