## Supplemental files for "CD271 restrains the B1b cell antibody response in a T cell dependent manner"

**Supplementary figure 1**

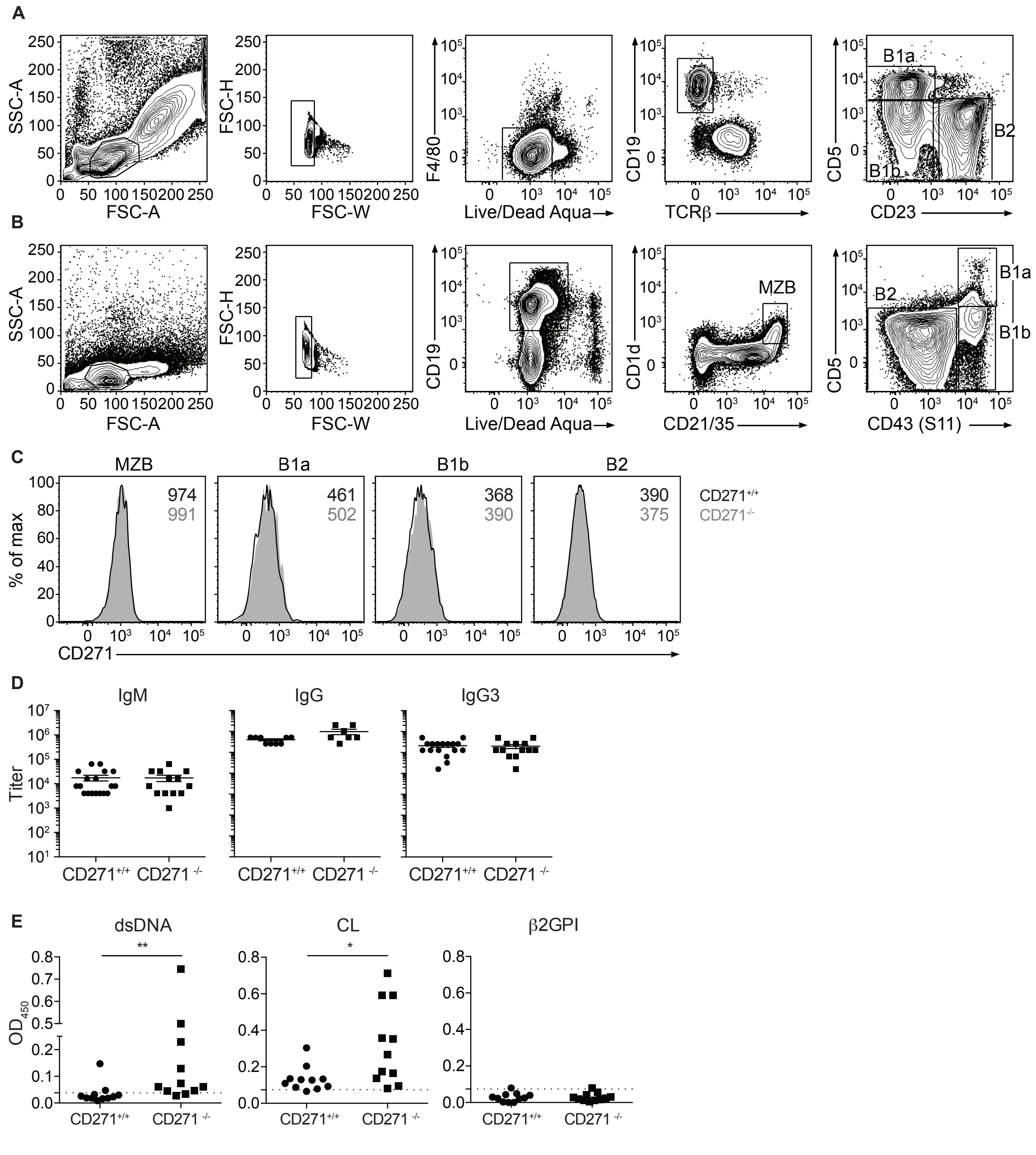


**Rare autoantibodies are detected in the serum of CD271-deficient mice.** Gating strategy of the B cell populations in the peritoneal cavity (**A**) and the spleen (**B**). (**C**) Representative histograms of CD271 expression on B cells isolated from the spleen of WT (black line) or CD271 KO mice (gray histogram). (**D**) Quantification of total IgM, IgG, and IgG3 in the serum of naive WT and CD271 KO mice. n=7-18 mice per group. (**E**) Serum from 1 year old WT or CD271-deficient naive male mice was harvested, and the level of double stranded DNA (dsDNA), cardiolipin (CL), or β2 glycoprotein I (β2GPI) autoantibodies were determined by ELISA. n=11 mice per group. Dots indicate individual mice. $*p < 0.05$, $**p < 0.01$

**Supplementary figure 2**

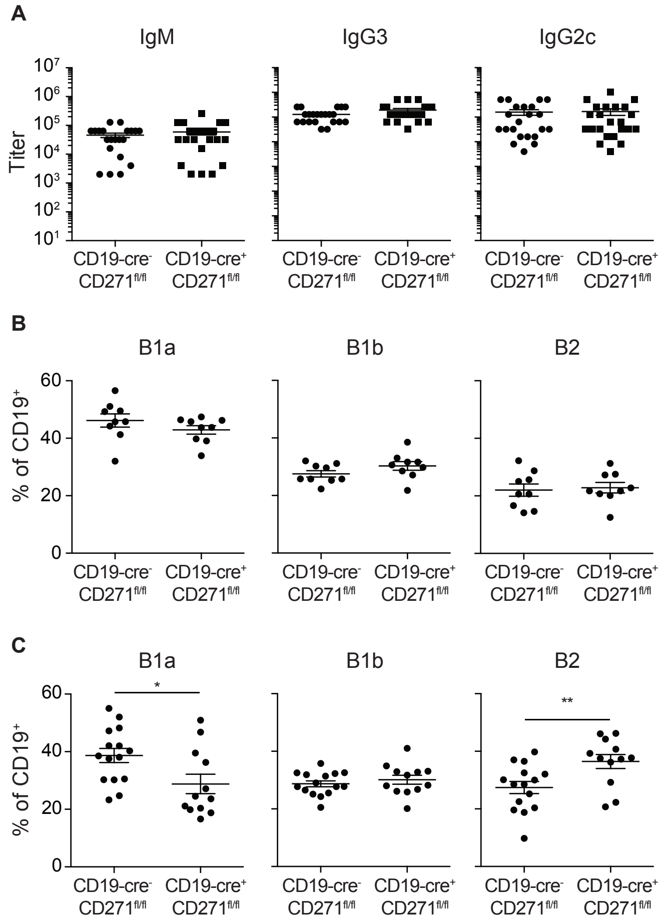


**Total serum antibodies and B1b cell proportions remain unchanged in the absence of B cell expressed CD271.** (**A**) Quantification of total IgM, IgG3 and IgG2c in the serum of naive CD19-Cre$^-$ CD271$^{fl/fl}$ and CD19-Cre$^+$CD271$^{fl/fl}$ mice. n=21-23 mice per group. Analysis of the proportion of the different B cell subtypes in the peritoneal cavity of CD19-Cre$^+$CD271$^{fl/fl}$ naive (**B**) and day 28 NP-Ficoll immunized mice (**C**) as compared to littermate controls (CD19-Cre$^-$CD271$^{fl/fl}$). n=9-15 per group. Dots indicate individual mice. *$p < 0.05$, **$p < 0.01$

**Supplementary figure 3**

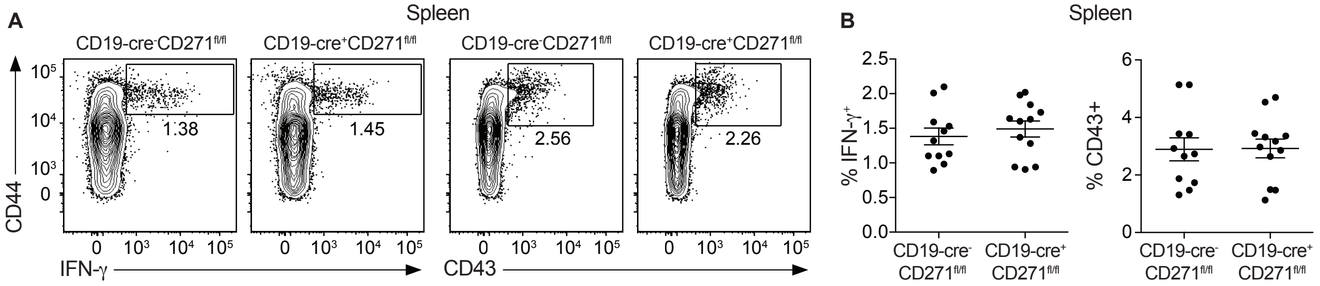


**T cell quiescence is unaffected in the spleen of CD271 KO mice.** $CD4^+$ T cells were isolated from the peritoneal cavity of naive CD19-$Cre^+$CD271$^{fl/fl}$ or littermate control (CD19-$Cre^-$CD271$^{fl/fl}$) mice and were stimulated with PMA/iono for 4 hours in the presence of Brefeldin A. Representative histograms (**A-C**) and compilation of the percentage (**D**) of IFN-$\gamma^+$, $CD43^+$, and $CD25^+$ $CD4^+$ T cells. n=11-12 mice per group. Dots indicate individual mice.
